# Symbiosis-driven development in an early branching metazoan

**DOI:** 10.1101/2022.07.21.500558

**Authors:** Aki H. Ohdera, Justin Darymple, Viridiana Avila-Magaña, Victoria Sharp, Kelly Watson, Mark McCauley, Bailey Steinworth, Erika M. Diaz-Almeyda, Sheila A. Kitchen, Angela Z. Poole, Anthony Bellantuono, Sajeet Haridas, Igor V. Grigoriev, Lea Goentoro, Elizabeth Vallen, David M. Baker, Todd C. LaJeunesse, Sandra Loesgen, Mark Q. Martindale, Matthew DeGennaro, William K. Fitt, Mónica Medina

## Abstract

Microbes can initiate developmental gene regulatory cascades in animals. The molecular mechanisms underlying microbe-induced animal development and the evolutionary steps to integrate microbial signals into regulatory programs remain poorly understood. In the upside-down jellyfish *Cassiopea xamachana*, a dinoflagellate endosymbiont initiates the life stage transition from the sessile polyp to the sexual medusa. We found that metabolic products derived from symbiont carotenoids may be important to initiate *C. xamachana* development, in addition to expression of conserved genes involved in medusa development of non-symbiotic jellyfish. We also revealed the transcription factor COUP is expressed during metamorphosis, potentially as a co-regulator of nuclear receptor RXR. These data suggest relatively few steps may be necessary to integrate symbiont signals into gene regulatory networks and cements the role of the symbiont as a key trigger for life history transition in *C. xamachana*.

## Main Text

The importance of microbes as regulators of animal physiology and life history is increasingly evident, from drivers of evolutionary adaptations with implications beyond individual organisms to supporting ecosystems, *e*.*g*., coral reefs (*1-4*). In particular, induction of developmental processes such as metamorphosis and organogenesis by microbes is common across metazoans, and may have facilitated the evolution of multicellularity in animals (*5, 6*). Despite the importance of microbes in shaping animal development and evolution, the molecular and evolutionary steps leading to the coupling of microbial signaling with animal developmental pathways remain largely unknown.

We used an emerging model system, the upside-down jellyfish *Cassiopea xamachana* (Cnidaria: Scyphozoa), to investigate how a photosynthetic dinoflagellate endosymbiont is integrated as a developmental cue to initiate metamorphosis (Fig. 1A). In non-symbiotic jellyfish, metamorphosis is triggered by environmental factors such as temperature (*7*) (Fig. 1B). For *C. xamachana*, the metamorphic transition in jellyfish known as strobilation begins approximately 10 to 17 days after acquisition of its endosymbiont *Symbiodinium microadriaticum* by the sessile polyp (scyphistoma) stage (Dinoflagellata: Symbiodiniaceae) (*8*). If symbionts are not acquired from the environment, *C. xamachana* will remain in the asexual scyphistoma stage indefinitely.

**Fig. 1.**
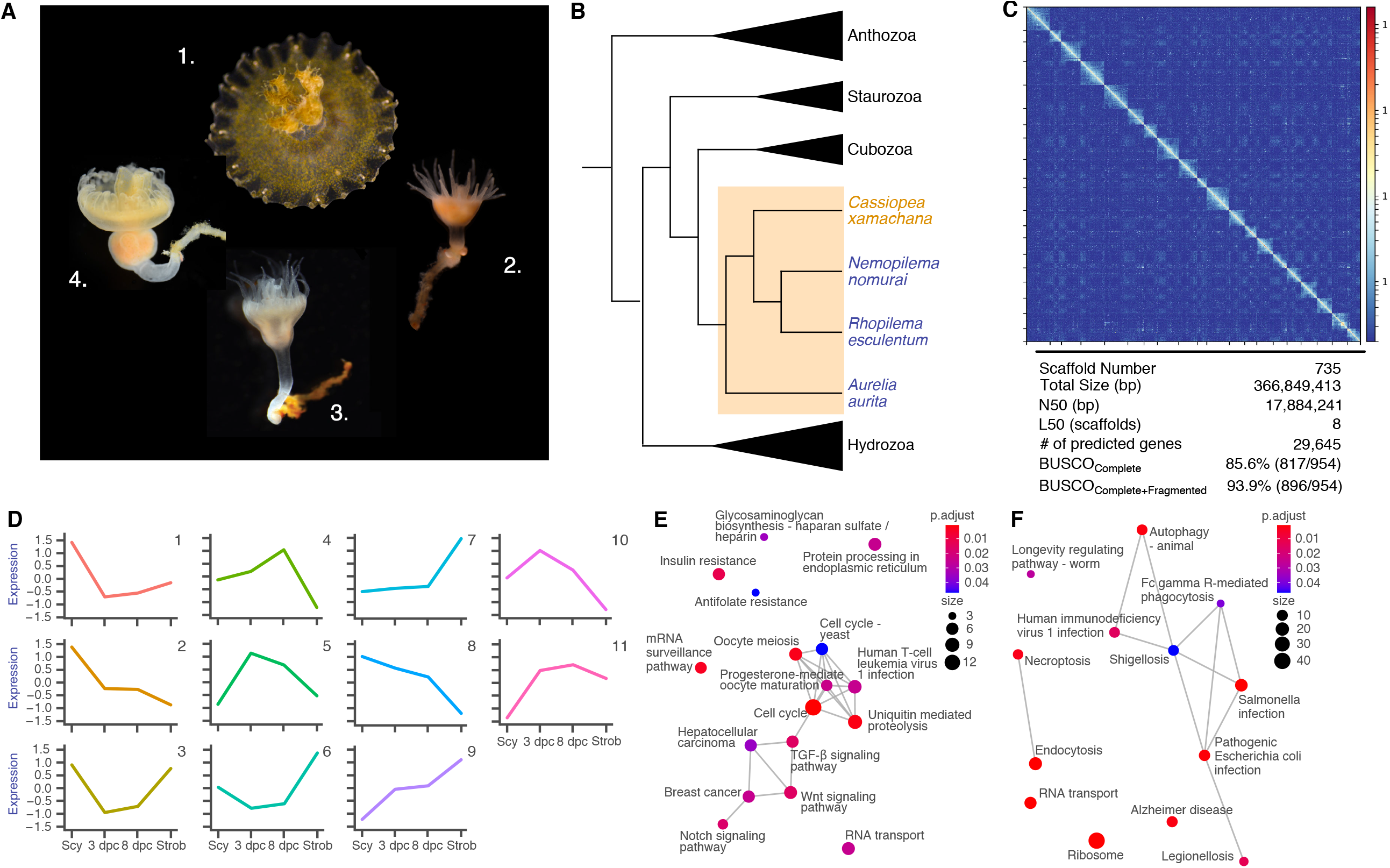
Symbiont induced metamorphosis coincides with activation of canonical developmental genes. (A) Lifecycle of *C. xamachana* displaying in clockwise order a free-swimming ephyra, aposymbiotic scyphistomae, symbiotic scyphistomae mid-strobilation (strobila), symbiotic scyphistomae during late strobilation. (B) Phylogenetic relationship of scyphozoan (colored box) jellyfish species relative to other cnidarian lineages. Species that metamorphosis under environmental factors are indicated in blue, while metamorphosis in *C. xamachana* (orange) is regulated by its symbiont. (C) HiC plot of the *C. xamachana* genome v2.0. Color bar indicates frequency of contact at each coordinate within the scaffolds. (D) Differentially expressed genes identified with ImpulseDE2 were clustered (k-means, k=11) according to co-expression patterns across the four experimental time points. Cluster centroids are plotted with standardized expression (E) KEGG pathways enriched in cluster 9 of k-means clustered genes exhibiting impulse-like expression. (F) KEGG pathways enriched in cluster 1 of k-means clustered genes exhibiting impulse-like expression.

We performed a differential mRNA expression analysis across time at 0, 3, and 8 days post-colonization (d.p.c) and mid-strobilation (∼ 17 d.p.c) of *C. xamachana* with ImpulseDE (*9*). We aligned the RNAseq reads to our new chromosome level *C. xamachana* genome assembly (NCBI accession OLMO00000000) composed of 20 pseudo-chromosomes (N_50_=17.9 Mb) that captured 99% of the original assembly (Fig. 1C), and contained 29,645 predicted protein coding genes (annotated genome available from https://phycocosm.jgi.doe.gov/Casxa1). We identified 5,414 genes (p-adjusted < 0.05) exhibiting an impulse-like (time-associated) expression pattern that grouped into 11 clusters (Fig. 1C). In clusters where expression increased with strobilation, genes involved in developmental regulation were enriched. Wnt, Ras, and cAMP signaling pathways were enriched in cluster 7, and Notch signaling and cell cycle pathways were enriched in cluster 9 (Fig. 1D,E; fig. S1, Supplementary File 1). Multiple pathways associated with animal-microbe interactions were also found in cluster 1 (626 DEGs, Fig. 1E, fig. S1), characterized by genes that remained stable after a downregulation at 3 d.p.c, and likely reflecting a transcriptomic response coinciding with the immune suppression characteristic of symbiosis maintenance in cnidarians (*10*).

We took advantage of the characterized molecular components of strobilation in non-symbiotic jellyfishes *Aurelia aurita* and *Rhopilema esculentum* to investigate their contribution to metamorphosis in *C. xamachana*. In non-symbiotic jellyfish, strobilation is regulated by an ortholog of *RXR* (*11, 12*), a nuclear receptor with a conserved function in metamorphosis across diverse metazoan lineages including vertebrates (*11-15*). In *A. aurita* and *R. esculentum*, expression of *RXR* is steadily up-regulated up to and through strobilation. Additionally, *retinol dehydrogenase* (*RDH*), a gene involved in the biosynthesis of the RXR ligand 9-cis retinoic acid (RA), also regulates metamorphosis in these non-symbiotic jellyfish (Fig. S2) (*11, 16*). In *C. xamachana*, we found *CxRXR* expression to be relatively stable across all time-points (Fig. 2A) and orthologs of *RDH1* and *RDH2* were either down-regulated 2-fold (*CxRADHa*, Fig. 3A) or did not show an impulse-like behavior (*CxRADHb*) compared to aposymbiotic scyphistomae. An additional gene, *CxCL112*, was recovered from a homology search of three hormone-like genes linked to strobilation in *A. aurita* (*CL112, CL390, CL631*; fig. S4) (*11*). In contrast to the gradual upregulation of *CL112* in *A. aurita, CxCL112* was down-regulated in the strobilating scyphistoma (strobila) stage relative to the aposymbiotic scyphistoma in *C. xamachana* (Fig. 2A, p-adjusted < 0.05). The only gene in which *C. xamachana* expression was similar to *A. aurita* during strobilation was of *DNA methyltransferase 1* (*DNMT1*; ∼ 1 log_2_FC, p-adj = 0.009; Fig. S3), consistent with a conserved role of this gene in development of animals (*17*). Thus, an initial interrogation of gene expression in *C. xamachana* during strobilation appears to show transcriptomic divergence between symbiotic and non-symbiotic species.

**Fig. 2.**
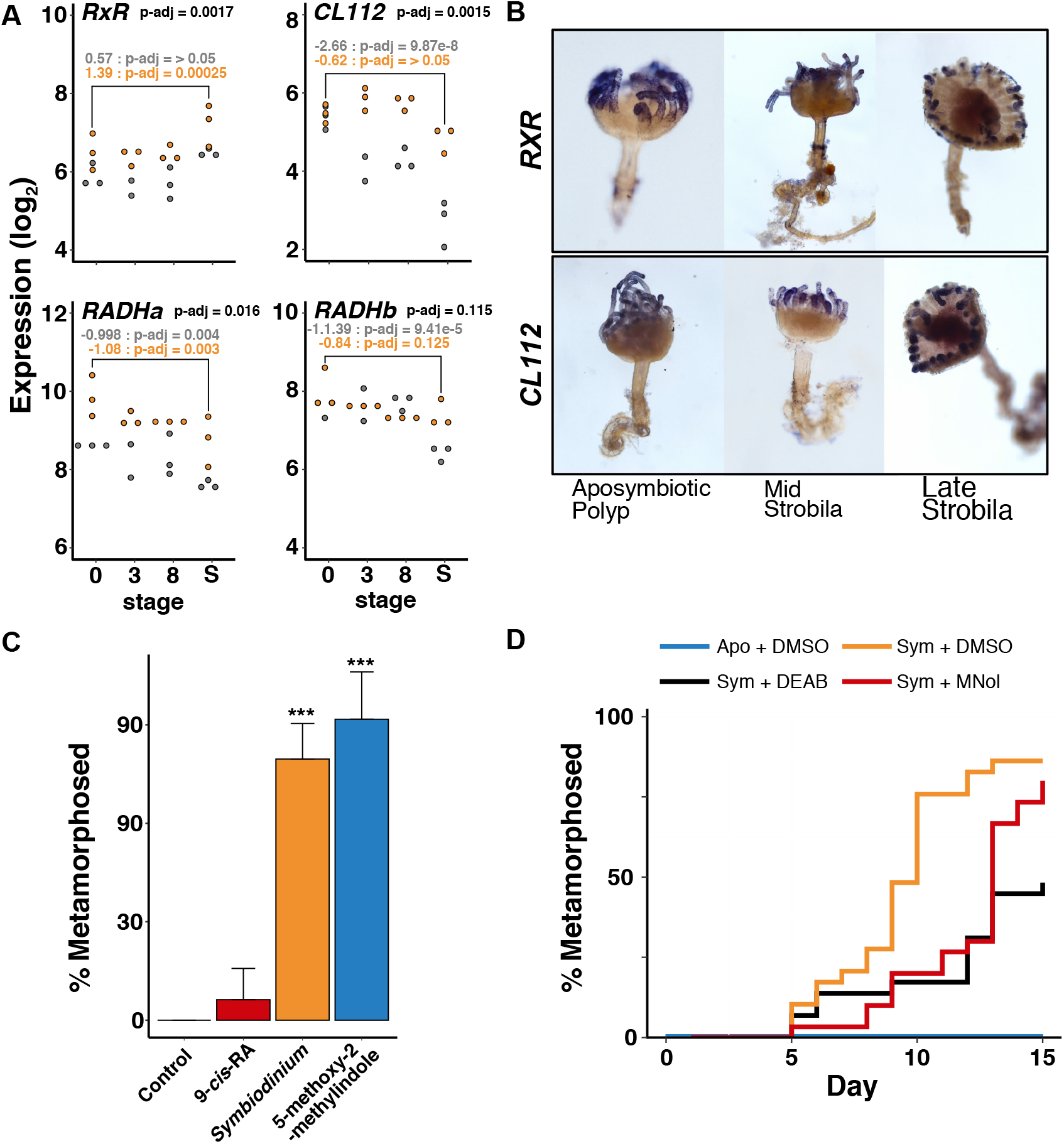
Regulation of retinoic acid pathway associated genes during strobilation in *Cassiopea xamachana*. A) Expression of genes implicated in non-symbiotic jellyfish strobilation across all stages (0=aposymbiotic scyphistomae, 3 and 8 days post-colonization, S=strobila). Values are log_2_ transformed DESeq2 normalized read counts. Colors indicate experiment 1 (orange) and experiment 2 (grey). P-adjusted values from the ImpulseDE analysis are indicated in black. P-adjusted values generated using DESeq2 are in orange (Exp. 1) and grey (Exp. 2). (B) Whole mount *in situ* hybridization of *CxRXR* and *CxCL112* in aposymbiotic (0 days) and strobila. Late-strobila were collected near completion of strobilation. (C) Strobilation rates of *C. xamachana* scyphistomae after 17 days post-treatment / colonization. ***p-value < 0.005 ANOVA with *post-hoc* Tukey test. (E) Strobilation rates of scyphistomae treated with an aldehyde dehydrogenase inhibitor (DEAB) and a COUP inhibitor (MNol) in symbiotic scyphistomae (n=40 / treatment group). DEAB (p-value < 0.0005, Mantel-Cox test) treated and MNol (p-value < 0.005, Mantel-Cox test) treated animals strobilated significantly slower than controls.

**Fig. 3.**
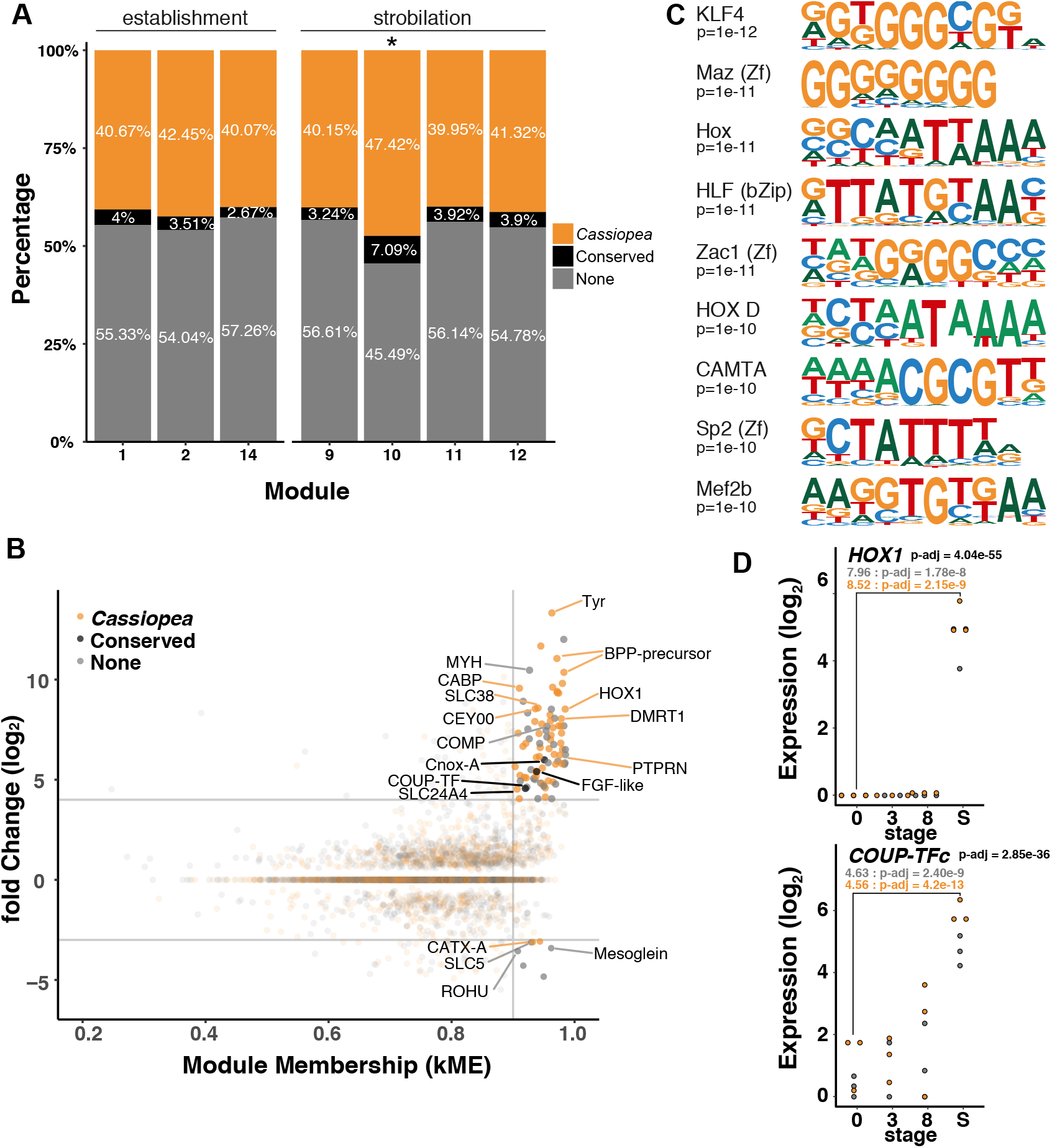
Developmental transcription factors are correlated with strobilation in *Cassiopea xamachana*. (A) Percentage of genes with an RXR response element (RE) within 5,000 bp of the gene. RXR-RE associated genes with orthologs in *A. aurita, N. nomurai*, and *R. esculentum* are shown in black. Non-conserved genes are shown in orange and genes lacking a proximal RXR-RE are shown in grey. Module 10 had a statistically significant number of genes compared to other modules (p-value = 5.70e^-6^ chi-squared test). (B) Module membership (kME) plotted against log_2_ fold-change. Dotted lines indicate (kME) cutoff of 0.9, and log_2_ fold-change cutoff (5, -3.5). (C) Position weight matrix logos for the top 10 most significantly enriched eukaryote motifs (p-value cutoff < 1 × 10^−10^) associated with genes in WGCNA modules correlating with strobilation (D) ImpulseDE2 normalized read counts of *Hox1* and *COUP-TF*c across the four time points, which showed a statistically significant impulse pattern during a time point.

When *CxRXR* and *CxCL112* mRNA were visualized by *in situ* hybridization, both genes were expressed orally in the aposymbiotic scyphistoma and mid-strobila stages, marking the region of the scyphistoma body that would undergo metamorphosis (Fig. 2B, fig. S5A,B). A decrease in expression of both genes coincided with progression of strobilation (late strobila), whereby mRNA became restricted to the retracted tentacles and the central oral appendage. Surprisingly, the diminished expression of *CxRXR* and *CxCL112* coinciding with completion of strobilation resembles the spatial pattern observed in *A. aurita* (fig. S5C). The expression data suggest much of the regulatory machinery identified in non-symbiotic jellyfish remains intact and may be important for *C. xamachana* development, albeit with several modifications.

We therefore tested whether treatment of aposymbiotic *C. xamachana* scyphistomae with exogenous chemicals that induce strobilation of non-symbiotic jellyfish (9-cis RA, 5-methoxy-2-methyl indole) would be as efficient in triggering metamorphosis compared to acquisition of symbionts (*16*). The potent strobilation inducer 5-methoxy-2-methylindole, a molecule containing a predicted pharmacophore of the *A. aurita* gene *CL390*, induced strobilation in *C. xamachana* at similar rates to *A. aurita* (91.7 %, one-way ANOVA with *post-hoc* Tukey test, p-value < 0.0005). While the target of the chemical inducer is not proposed hormone gene *CL390*, both *A. aurita* and *C. xamachana* share an inducible strobilation regulator. Interestingly, 9-cis RA at saturation (1 uM) failed to significantly induce strobilation relative to the vehicle control (6.25 %, p-value > 0.05, Fig. 2C). We thus hypothesized that strobilation in *C. xamachana* may require additional regulators and activation of RXR by 9-cis RA is insufficient to induce complete strobilation.

To identify genes correlated with strobilation and symbiosis establishment, we next performed weighted gene correlation network analysis (WGCNA) (*18*). We identified three modules (2,846 genes, Modules 1, 2, and 14) that were significantly correlated with symbiosis establishment (3 and 8 d.p.c), and four modules (3,714 genes, Modules 9,10,11, and 12) associated with strobilation (p-value < 0.05) (fig. S6A, Supplementary File 2). Genes in the strobilation modules were enriched in pathways responsible for energy modulation and metabolism (mTOR signaling, insulin signaling, glucagon signaling, oxidative phosphorylation) coinciding with developmental processes, as well as cell proliferation and morphogenesis (cell cycle, DNA replication, TGF-β signaling) (fig. S6B).

We investigated whether genes in the strobilation module were associated with the conserved *RXR* response element (AGGTCA). If *RXR* regulates strobilation in *C. xamachana*, we expect the response element to be enriched with genes linked to strobilation. We scanned the 5,000 bp region flanking each gene for RXR half-sites separated by 1 to 5 nucleotides and found 1,676 genes of the strobilation module were proximal to a putative *RXR* response element (Supplementary File 3). Of these, we found 83 genes with orthologs in related jellyfish species with conserved proximity to an *RXR* response element (Supplementary File 4). Moreover, the strobilation module M10 was enriched with genes associated with an *RXR* response element (Fig. 3A, p-value < 5.70e^-6^ Chi-squared test). We next compared module membership (kME), high values akin to high inter-connectedness within the network, to fold-change. It revealed several “hub” genes including two *hox* genes and a chicken ovalbumin upstream promoter transcription factor (*COUP-TF*, nuclear receptor subfamily II) among those with high kME and expression (Fig. 3B, fig. S6A). Developmental genes including *Doublesex and Mab3-related* (*DMRT*), which is associated with strobilation in non-symbiotic jellyfish, were also among those with high kME values (*11*). Genomic scans for additional motifs that were significantly enriched (p-value < 1e^-10^) within the strobilation modules included binding targets of homeobox genes (Fig. 3C). Although with marginally lower significance (p-value < 1e^-5^), *COUP* motifs were also identified to be enriched (Supplementary File *5*), making them likely candidates to regulate strobilation in *C. xamachana*.

In further exploring *COUP-TF* we identified *CxCOUP-TFa*, which exhibited a stepwise pattern of expression that increased during days 3 and 8 d.p.c., and *CxCOUP-TFb*, which showed gradual up-regulation over time (fig. S7). A third closely related nuclear receptor (*CxCOUP-*like) showed a rapid increase in expression during strobilation, similar to patterns observed for other important players in strobilation (Fig. 3D). Members of the *COUP-TF*s exhibit dual roles as both initiators and repressors of transcription, with potential regulatory roles in embryogenesis through heterodimerization with the *RXR* gene (*19, 20*). Although it remains unknown whether *CxRXR* forms a heterodimer with other nuclear receptors in *C. xamachana*, the low induction of strobilation under 9-cis RA treatment suggests *CxRXR* activation of strobilation requires a co-regulator (*21*). To test the participation of a *COUP-TF* in *C. xamachana* strobilation, we treated scyphistomae harboring *S. microadriaticum* with a chemical inhibitor of COUP transcription factors, 4-methoxy-1-napthol (MNol). We found strobilation to be significantly delayed (Fig 2D), thus implicating *COUP-TF* as a key regulator of strobilation.

As carotenoids are common ligands of nuclear receptors like RXR we further investigated our datasets for genes involved in carotenoid metabolism. We found a β*-carotene oxygenase (CxBCOa*) up-regulated approximately four-fold during strobilation (p-value = 6.27e^-5^, Fig. 4A, fig. S6). BCO genes perform symmetric or asymmetric cleavage of β-carotene. β-carotene monooxygenase (BCMO) produces retinal, while β-carotene dioxygenase (BCDO2) generates β-ionone and β-apo-carotenal. These molecules can be further processed by a dehydrogenase to produce potential nuclear receptor ligands, *i*.*e*. retinoic acid (*22, 23*) (Fig. 4B). Two BCO-like genes (*CxBCOLb, CxBCOLf*) were also found in the WGCNA symbiosis establishment modules (M1, M14) (Fig. 4A, fig. S8). While the exact type of reaction performed by *CxBCOa* requires further characterization, key residues responsible for catalytic activity were found in all genes (fig. S9), suggesting cleavage activity is likely analogous with BCO orthologs in mammals. In mammals, BCDO2 is thought to be predominantly active in the mitochondria where it functions against oxidative damage caused by carotenoid accumulation, whereas BCMO is restricted to the cytosol (*24, 25*). None of the *CxBCO* genes possessed a mitochondrial transport signal within the N-terminus. However, signaling peptide sequences were present in *CxBCOL* genes, suggestive of their post-secretion enzymatic activity within the extracellular matrix (fig. S10) (*26*). Treatment of symbiotic scyphistomae with the RDH inhibitor 4-diethylaminobenzaldehyde (DEAB) led to significant delays to strobilation (Mantel-Cox test, p-value < 0.005) (Fig 2D). Treatment with a single concentration of butylated hydroxytoluene (BHT), a BCO inhibitor, did not alter rates of strobilation, but further experiments with additional concentrations may be useful (fig. S11). These results suggest carotenoids are important for strobilation in *C. xamachana*, and potentially sourced from the symbionts.

**Fig. 4.**
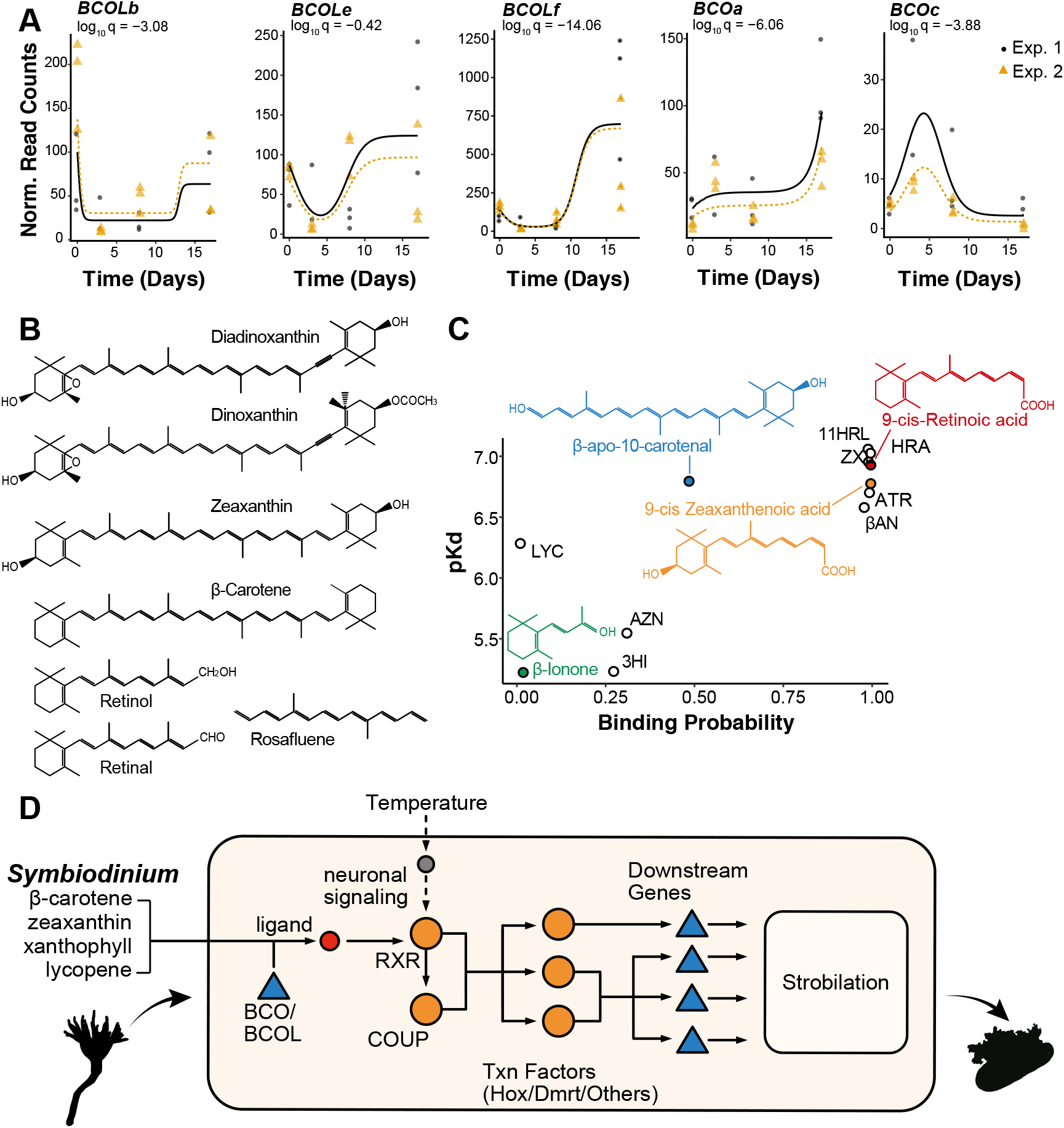
Potential RXR ligand interactions underlying regulation of symbiosis driven strobilation. (A) DESeq2 normalized read counts of *C. xamachana* β-carotene oxygenases found to be correlated with establishment / strobilation over the four time points. (B) Carotenoids produced by the dinoflagellate symbiont *Symbiodinium microadriaticum*. Retinol, retinal, and rosafluene are cleavage products of β-carotene. (C) Binding probability of potential ligands against their dissociation constant modeled with BindScope and Kdeep. 3HI = 3 hydroxy-beta-ionone. 11HRL = 11-cis-3-hydroxyretinal. ATR = all-trans retinoic acid. AZN = apo-9 zeaxanthenone. BAC = beta-10-apocarotenal. BAN = 13-apo-beta-apocarotenone. HRA=9-cis 4hydroxyretinoic acid. LYC=lycopene. ZXL=all-trans zeaxanthenal. (D) Proposed model of *C. xamachana* strobilation. Carotenoids produced by *Symbiodinium* are cleaved by *CxBCO*. Along with a secondary nuclear receptor (*e*.*g*., *COUP-TF*), *CxRXR* binds to response elements proximal to developmental genes (e.g., *Hox, DMRT*) to initiate the downstream developmental cascade leading to strobilation.

*Symbiodinium* synthesizes multiple carotenoid products that can be potential substrates for further processing (*27*). Genes required for carotenoid synthesis were confirmed to be expressed by *S. microadriaticum in hospite*, including those responsible for production of β-carotene, lycopene, peridinin, zeaxanthin, and dinoflagellate specific carotenoids (dinoxanthin, diadinoxanthin) (Fig. 4B, fig. S12A) (*28*). *In silico* predictions of binding affinities for putative carotenoid ligands suggests cleavage products other than 9-*cis* retinoic acid have similar binding kinetics with CxRXR, including zeaxanthenoic acid, a metabolic equivalent of RA derived from zeaxanthin (Fig. 4C, fig. S12B).

Taken together, our data suggest that the regulation of development in a symbiotic jellyfish remains largely similar to non-symbiotic species with some notable exceptions. We hypothesize that after colonization of *C. xamachana* scyphistomae by *S. microadriaticum*, accumulated carotenoids are processed by CxBCO, leading to the production of nuclear receptor ligands. Subsequently, nuclear receptors act as a sensor by binding these carotenoid cleavage products to trigger a downstream signaling cascade leading to the initiation of strobilation (Fig. 4E). With the exception of diadinoxanthin and peridinin, which generally comprise over 80% of the produced carotenoids in *Symbiodinium*, minor pigments make up less than 5% of the total (*29*). Consequently, the observed lag period to the start of strobilation following initial acquisition of symbionts may reflect the time necessary to accumulate the minimum concentration of a minor carotenoid (e.g., β-carotene, zeaxanthin) required to activate RXR and irreversibly initiate development. The difference in expression of key genes responsible for RA metabolism between *C. xamachana* and non-symbiotic jellyfish, including the constitutive expression of *RXR*, may also accelerate the time to strobilation by priming the animal for the transition. We suggest the integration of *Symbiodinium* as driver of development in *C. xamachana* likely resulted from a compatibility of precursor molecules of symbiont origin with the RXR developmental signaling pathway, accompanied by the modulation in expression of key genes. Although specific pathways may vary, symbiont-driven development may evolve through relatively few changes to existing programs.

## Acknowledgement

The work conducted by the U.S. Department of Energy Joint Genome Institute, a DOE Office of Science User Facility, is supported by the Office of Science of the U.S. Department of Energy under Contract No. DE-AC02-05CH11231.

## Notes

### Competing Interest Statement

The authors have declared no competing interest.

