## Supplemental File for "Symbiosis-driven development in an early branching metazoan"

### Materials and Methods

#### Animal Cultures

Apo-symbiotic polyps were maintained in 6-well culture plates at 26°C in the dark and fed 24 hr-old *Artemia* 2-3 times per week. Partial water changes were performed 24 hr after feeding, with care taken to remove unconsumed *Artemia*. *Symbiodinium microadriaticum* KB8 cultures were maintained with a 12:12 cycle at 60-100 $\mu E \cdot m^{-2} \cdot s^{-1}$ at 26°C.

#### Artificial and Natural Induction of Strobilation in *Cassiopea xamachana*

Stock solutions of 1 mM retinoic acid and 50 mM 5-methoxy-2-methylindole were made in DMSO. Solutions were added to 2 ml of 0.2 µM filtered artificial seawater in a 24-well plate for a final concentration of 1 µM and 50 µM of 9-cis retinoic acid and 5-methoxy-2-methylindole, respectively. A single polyp (N=16) was added to each well, and strobilation was monitored over a course of 5 days at 26°C. The well water was replaced daily with sea water freshly treated with chemicals. Strobilation rates in response to *Symbiodinium* were monitored over a course of 21 days. Polyps were exposed to 100,000 *Symbiodinium* cells with simultaneous feeding of *Artemia* for 24 hr and transferred to individual wells with 0.2 µm filtered artificial seawater. Polyps were maintained at 26°C at 100 $\mu E \cdot m^{-2} \cdot s^{-1}$ of 12:12 light-dark cycle and fed twice per week for the duration of the experiment. Strobilation was determined as complete upon release of the ephyra from the polyp.

#### RNAseq of colonization and strobilation

Aposymbiotic scyphistomae (polyps) from 6 clonal lines (T1-A, T1-B, T2-B, T1-E, T1-F, T2-E) were colonized with *Symbiodinium microadriaticum* (KB8). *Symbiodinium* cells were provided to *Artemia* at a concentration of 8.025 x $10^6$ cells prior to being fed to polyps. Polyps were allowed to feed for 2 hrs prior to being transferred to clean filtered artificial seawater. Polyps were maintained at 26°C with 100 $\mu E \cdot m^{-2} \cdot s^{-1}$ 12:12 light-dark cycle, with feeding every other day. A subset (minimum 15 polyps) was collected on each sampling day and immediately preserved in RNAlater, and stored in a -80ºC ultrafreezer for subsequent RNA extraction. For apo-symbiotic polyps, a set of 50 polyps was used to isolate enough total RNA for downstream processing. Polyps were collected at 3 days and 8 days post-colonization as well as during mid-strobilation when the tentacles have been resorbed and the calyx has shown evidence of undergoing constriction. Strobilae with pulsating ephyra were not sampled.

RNA was extracted using CTAB phenol-chloroform protocol. Polyps were homogenized with a bead beater using 250 µl of 0.1 and 0.5 mm zirconia beads in Qiazol. Chloroform was added directly to the bead tube prior to proceeding through the extraction protocol. RNA concentration was determined using Qubit 2.0 (Thermo Fisher Scientific) and quality assessed using the Bioanalyzer (Agilent) with the RNA 6000 Nano kit and NanoDrop 1000 spectrophotometer (Thermo Fisher Scientific). Extracted total RNA was preserved at -80°C until further processing. The mRNA was isolated from total RNA using the Dynabeads mRNA purification kit (Ambion). Sequencing libraries were constructed following the Joint Genome Institute's (JGI) custom Illumina stranded Truseq library protocol with an insert size of 250 bp. Sequencing was done in two batches. Libraries for T1-A, T1-B, and T2-B were sequenced on the Illumina HiSeq2000 to generate 150 bp single-end reads at the Pennsylvania State University Genomics Core Facility. T1-E, T1-F, and T2-E libraries were sequenced on the Illumina HiSeq2000 to generate 150 bp paired-end reads at JGI.

**Genome assembly**
The *Cassiopea xamachana* draft genome (version 1) (*1*) was re-assembled by Dovetail.

Chicago library preparation and sequencing
A Chicago library was prepared as described previously (*2*). Briefly, ~500 ng of HMW gDNA (mean fragment length = 50 kbp) was reconstituted into chromatin *in vitro* and fixed with formaldehyde. Fixed chromatin was digested with DpnII, the 5' overhangs filled in with biotinylated nucleotides, and then free blunt ends were ligated. After ligation, crosslinks were reversed and the DNA purified from protein. Purified DNA was treated to remove biotin that was not internal to ligated fragments. The DNA was then sheared to ~350 bp mean fragment size and sequencing libraries were generated using NEBNext Ultra enzymes and Illumina-compatible adapters. Biotin-containing fragments were isolated using streptavidin beads before PCR enrichment of each library. The libraries were sequenced on an Illumina HiSeq X to produce 213 million 2x150 bp paired end reads, which provided 48.72 x physical coverage of the genome (1-100 kb pairs).

Dovetail HiC library preparation and sequencing
A Dovetail HiC library was prepared in a similar manner as described previously (*3*). Briefly, for each library, chromatin was fixed in place with formaldehyde in the nucleus and then extracted Fixed chromatin was digested with DpnII, the 5' overhangs filled in with biotinylated nucleotides, and then free blunt ends were ligated. After ligation, crosslinks were reversed and the DNA purified from protein. Purified DNA was treated to remove biotin that was not internal to ligated fragments. The DNA was then sheared to ~350 bp mean fragment size and sequencing libraries were generated using NEBNext Ultra enzymes and Illumina-compatible adapters. Biotin-containing fragments were isolated using streptavidin beads before PCR enrichment of each library. The libraries were sequenced on an Illumina HiSeq X to produce 249 million 2x150 bp paired end reads, which provided 4,776.27 x physical coverage of the genome (10-10,000 kb pairs).

Scaffolding the assembly with HiRise
The input *de novo* assembly, shotgun reads, Chicago library reads, and Dovetail HiC library reads were used as input data for HiRise, a software pipeline designed specifically for using proximity ligation data to scaffold genome assemblies (Putnam et al, 2016). An iterative analysis was conducted. First, Shotgun and Chicago library sequences were aligned to the draft input assembly using a modified SNAP read mapper (http://snap.cs.berkeley.edu). The separations of Chicago read pairs mapped within draft scaffolds were analyzed by HiRise to produce a likelihood model for genomic distance between read pairs, and the model was used to identify and break putative misjoins, to score prospective joins, and make joins above a threshold. After aligning and scaffolding the Chicago data, Dovetail HiC library sequences were aligned and scaffolded following the same method. After scaffolding, shotgun sequences were used to close gaps between contigs.

Scaffolds potentially containing bacterial genes were removed from the draft genome assembly (version 1) and the dovetail assembly (version 2) using EukRep (*4*). Removed scaffolds were manually screened, and those which did not contain known bacterial genes were retained in the final assembly. The final assembly was composed of 735 scaffolds, with an N50 of 17.6 Mb and a predicted genome size of 366 Mb. Approximatey 99% of the genome was captured within the 20 largest scaffolds. We used HiCExplorer to generate and visualize the Hi-C contact matrix (*5*). Genome completeness was determined using BUSCO against the metazoa reference set v? (*6*).

**Gene Model Predictions**

Repeat families were identified using RepeatModeler v1.0.11 (https://github.com/Dfam-consortium/RepeatModeler) into a custom library used to mask the genome assembly with RepeatMasker v4.0.7 (http://www.repeatmasker.org/RepeatModeler/). *Ab initio* gene prediction was performed using GeneMark-ES v4.38 with the following parameters: -- soft_mask 5000, --max_intron 25000 (*7*). The BRAKER2 pipeline was used for RNAseq-informed gene prediction (*8*). RNA paired-end reads were aligned to the EukRep filtered genome using HiSAT2 v2.1 (*9*). HiSAT2 read mapping was performed with the following parameters: --mp 1,0 --pen-noncansplice 20. Genes predicted with BRAKER2 with less than 90% read coverage over the exon boundaries were removed and reads retained were classified as high-quality. PASA v2.3.3 was used to predict genes using a *de novo* assembled transcriptome (*10*). RNAseq reads mapped with HiSAT2 were assembled using Trinity v2.8.5 with default settings (*11*). Protein evidence was generated using Exonerate v2.2.0 with the following settings: --percent 80, --ryo "AveragePercentIdentity: %pi\n" against all Cnidaria genes in the Uniprot database (*12, 13*). All gene predictions were combined with EVidenceModeler v1.1.1 with the following weights: PROTEIN exonerate 1, TRANSCRIPT pasa 10, OTHER_PREDICTION HiQuality 5, ABINITIO_PREDICTION Augustus 1, ABINITIO_PREDICTION GeneMark 1 (*14*). An additional round of PASA prediction was performed to predict UTRs. We predicted 29,645 genes and 2,893 isoforms (33,538 genes total).

Genes were annotated as the top matches from blastx searches against the nr and Swissprot database (*15, 16*). Multiple sequence alignments of short-chain dehydrogenases and β-carotene oxygenases were obtained from MAFFT v7.4 (gap extend penalty = 0.123, gap opening penalty = 1.53, and matrix = BLOSUM62) through the ngPhylogeny web interface (*17, 18*). The alignment was trimmed for non-informative sites using BGME and a maximum likelihood tree was constructed using PhyML v3.1 with automatic model selection and bootstrapping (n=500) (*19*). Trees were visualized using the interactive Tree of Life (iTOL) v4 (*20*).

Similar approaches were taken for the nuclear receptor superfamily. Representative sequences of nuclear receptors from mammals and *Drosophila melanogaster* were retrieved from NCBI or Uniprot, and BLAST against the *C. xamachana* protein models. NR sequences from *C. xamachana* were appended to the original query set, and BLAST was repeated against protein models of *Tripedalia* and *Aurelia aurita*. The protein family tree was constructed using the methods described above. Sequences were manually curated after initial MAFFT alignment, prior to trimming with BGME.

**RNAseq Analysis**

Trimming and adapter removal was done with Trimmomatic-0.36 (Bolger et al., 2014). and genome-guided assembly was conducted using Trinity v2.4.0 using standard parameters (*11, 21*). Reads were aligned to the genome containing using STAR with the following parameters to optimize alignment: --limitOutSJcollapsed 100.1200000 --limitSjdbInsertNsj 1000000 --outFilterMultimapNmax 100 --outFilterMismatchNmax 33 --outFilterMismatchNoverLmax 0.3 --seedSearchStartLmax 12 --alignSJoverhangMin 15 --alignEndsType Local --outFilterMatchNminOverLread 0 --outFilterScoreMinOverLread 0.3 --winAnchorMultimapNmax 50 --alignSJDBoverhangMin 3. Differential expression analysis of the host and symbiont was conducted using DESeq2 on both batches of sequencing, separately (*22*) (Fig. S12A,B). Differentially expressed genes for each time point were identified via pairwise contrasts between the apo-symbiotic state (apo v 3/8 days, apo v strobila) (Supplementary File 6).

In order to analyze the batches simultaneously as a single dataset across multiple time points, we used ImpulseDE2 (*23*) (Supplementary File 2). Batch correction was applied and impulse-like behavior was identified. WGCNA analysis was performed across both batches. Sample T1-Bt3 was removed as it did not cluster with other replicates. Reads were initially normalized with DESeq2, and adjusted for batch effect using the limma v3.40.6 package in R (*24*). The soft-threshold power ® was set to 28 to calculate network adjacency, which was further transformed into a topological overlap matrix (TOM) to reduce noise and calculate the dissimilarity. Initially identified modules containing genes with similar expression profiles were merged and analyzed for trait associations (symbiotic-state, establishment, strobilation) (Supplementary File 4). KEGG enrichment analysis of genes associated with establishment and strobilation was performed with the *enrichKEGG* function of the clusterProfiler R-package, with the KEGG Orthology database as the background (*25*) (Supplementary Files 2,5).

**Whole genome alignments and ortholog assignment**

Whole genome alignment of *Aurelia aurita* (NCBI Accession: GCA_004194415.1), *Rhopilema esculentum* (Accession: GCA_013076305.1), and *Nemopilema nomurai* (GCA_003864495.1) were conducted with Cactus v1.1.1 (*26*). In order to identify orthologs, duplicate alignments were filtered and only alignments in which sequences from all four taxa were included were retained. With *C. xamachana* as reference, coordinates for each sequence alignment were lifted from the target species, and coordinates intersecting with the gene models were extracted using bedtools v2.25.0. We also performed pairwise reciprocal BLAST between the *C. xamachana* protein models and each other scyphozoa dataset to confirm orthologs identified with the Cactus algorithm.

**Transcription factor Motif Analysis**

Motif search for putative RXR binding sites within the 5,000 bp flanking the predicted transcript models was performed using HOMER v4.11 (*27*). We searched for direct repeats with positional weight matrices consisting of two repeating AGGTCA motifs with 1 – 5 nucleotide spacers with a variable site tolerance of two. Scans for *de novo* and known motifs proximal (5,000 bp) to genes within the co-expressed modules identified with WGCNA were also performed with HOMER, using the 5,000 bp flanking region of all other genes as background. The difference in the proportion of RXR binding sites identified for each co-expression module was compared using a Pearson's Chi-square test in R.

***In Situ* Hybridization**

Adult gonadal tissue was used to isolate mRNA using the RNA extraction method described above (see "RNAseq Analysis"). cDNA was synthesized using Advantage RT for PCR kit (Clontech). Primers were designed for *CxRXR*, *CxCL112*, and *CxMcolA* (positive control) with a target amplicon length of greater than 1000 bp. Amplified target genes were ligated into plasmids using the pGEM-T cloning kit (Promega) cloned into Dh-5α competent *E. coli* cells. Successful cloning was confirmed via colony PCR and plasmids were extracted using the Hi-Speed Mini Plasmid Kit (IBI). Plasmids were Sanger sequenced to ensure orientation of insertion of target sequences into the plasmid. Target sequences were amplified using T7 and SP6 primers and purified from a gel using the QiaQuick extraction kit (Qiagen). Final riboprobes were synthesized using the Megascript Sp6 and T7 Transcription kit (Thermo Fisher Scientific) to generate both sense and anti-sense mRNA. Riboprobes were stored in hybridization buffer (50% Formamide, 10% 20X SSC pH 4.5, 0.25% 20mg/ml heparin, 0.25% Tween-20, 1% of 20% SDS) with a final concentration of 100 ug/ml salmon sperm DNA. Riboprones were stored at -80°C until further use.

Whole mount *in situ* hybridization was performed with polyps collected immediately prior to the start of a slightly modified protocol described by Wolenski et al. (2013). Polyps were incubated in proteinase K for 25 minutes at a final concentration of 0.01 mg/ml. Samples were incubated in the pre-hybridization buffer overnight at 60°C. Both sense and anti-sense probes for each target were used to control for non-specific hybridization. Samples were hybridized in each probe at a concentration of 1 ng/ml for 48 hrs at 60°C. Blocking of samples was carried out at 4°C for 1 hr prior to an overnight incubation in the anti-DIG, AP Fab fragments at 4°C. For mRNA detection, samples were developed in alkaline phosphatase substrate solution for 15 min to 1 hr prior to halting the reaction. Processed samples were washed with PBS-tween and mounted in 80% glycerol for imaging and long-term storage at 4°C.

**Pharmacological inhibition of strobilation**

Aposymbiotic *C. xamachana* polyps were incubated with 5 x $10^5$ *S. microadriaticum* KB8 cells at a density of 10,000 cells/ml artificial seawater for three days prior to the start of the experiment, reserving 60 aposymbiotic polyps for negative control (i.e., no symbiont exposure) groups. Polyps were maintained under a 12:12 cycle at 60-100 $\mu E\square m^{-2}\square s^{-1}$ at 26°C. After the three day incubation, polyps were randomly assigned a treatment and sorted into a single well of a 48-well culture plate. 30-31 animals were assigned to each treatment in total. Colonized polyps were treated with 10 µM 2,6-Di-tert-butyl-4-methlphenol (BHT, N=30), 3 µM 4-methoxy-1-napthol (MNol, N=30), 50 µM diethylaminobenzaldehyde (DEAB, N=31). Polyps were also treated with the RXR inhibitor UVI3003 at a concentration of 1 µM (N=31). These concentrations were based on previous literature demonstrating inhibition by the molecules on their respective pathways or, in the cases of Mnol and UVI3003, were otherwise established by titrating down to doses that did not cause obvious signs of physiological stress. For the case of UVI3003, concentrations above 1 µM were toxic and only survived dilute dosages, resulting in no effect on strobilation rate. Four vehicle control groups were included with the experiment: aposymbiotic polyps treated with DMSO, aposymbiotic polyps treated with ethanol, symbiotic polyps treated with DMSO, and symbiotic polyps treated with ethanol (N=30 in each case). Polyps were fed *Artemia* twice a week during the duration of the experiment, and moved into clean artificial seawater with freshly prepared chemicals 24 hrs after each feeding. Strobilation Strobilation was assessed every day and scored at the first unambiguous sign of metamorphosis (typically changes in the shape of the margin around the mouth of the polyp); this differs from the method described in "Induction of Strobilation" because the various inhibitors may differentially hinder progression through the steps of metamorphosis.

***In silico* ligand binding prediction**

The protein structure of *C. xamachana RXR* was modeled based on homology using SWISS-MODEL via the ExPasy web server (*28, 29*). The binding pocket coordinates were predicted using DeepSite (*30*). Protein-ligand binding was predicted with BindScope, employing a threedimensional convolutional neural-network approach (*31*). The 3D ligand structures in sdf format were downloaded from PubChem or created using molView.org (Supplementary File 6). The output of BindScope was used to predict the binding affinity of each ligand utilizing a deep convolutional neural networks approach with Bin $K_{DEEP}$ (*32*).

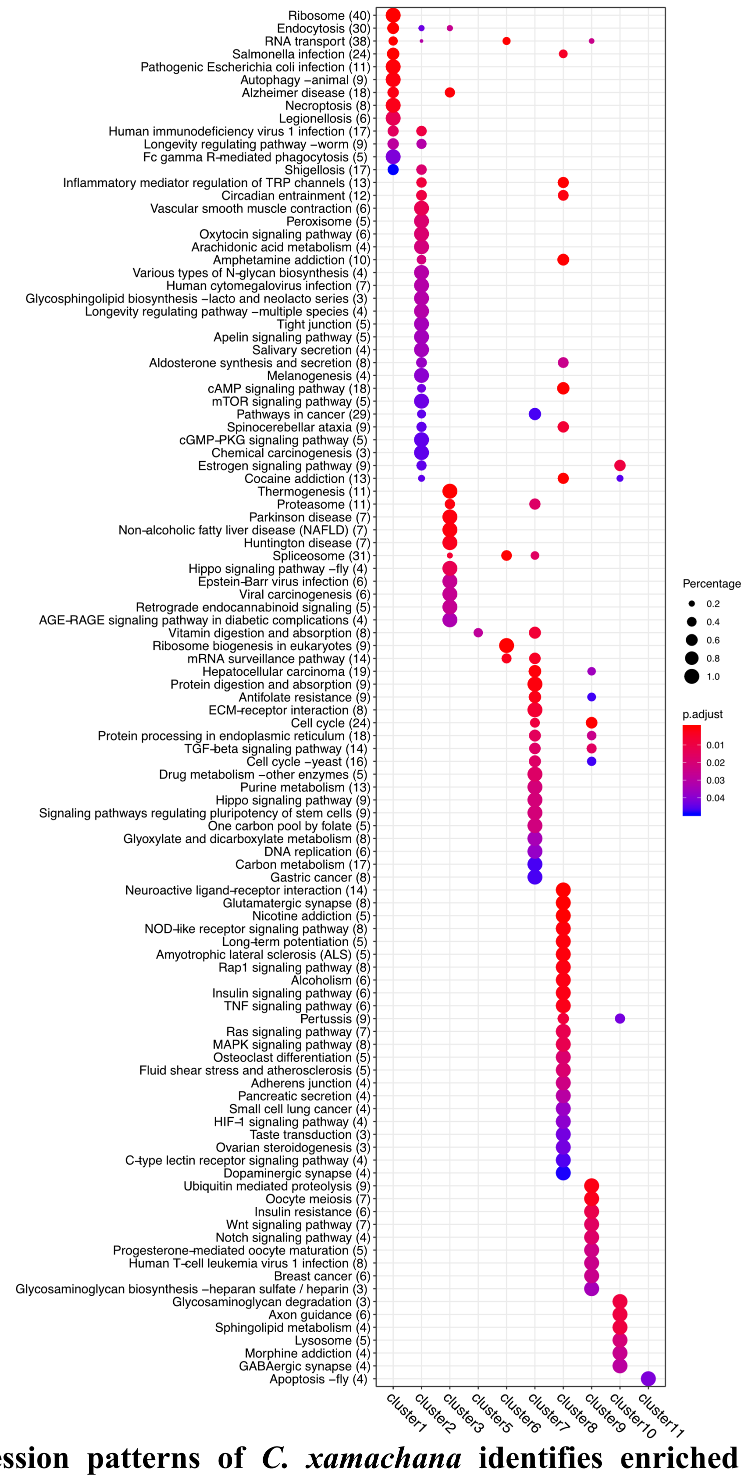


**Figure S1: Expression patterns of *C. xamachana* identifies enriched KEGG pathways capturing processes in development and onset of symbiosis.**

KEGG enrichment analysis with ClusterProfiler captures genes involved in development (cluster 9) and symbiosis (cluster 1) that reflect predicted expression patterns. Percent recovery of genes encompassed by the KEGG pathways are indicated by the size of the circle. Enrichment score (p-adjusted) is indicated by the color of the circle.

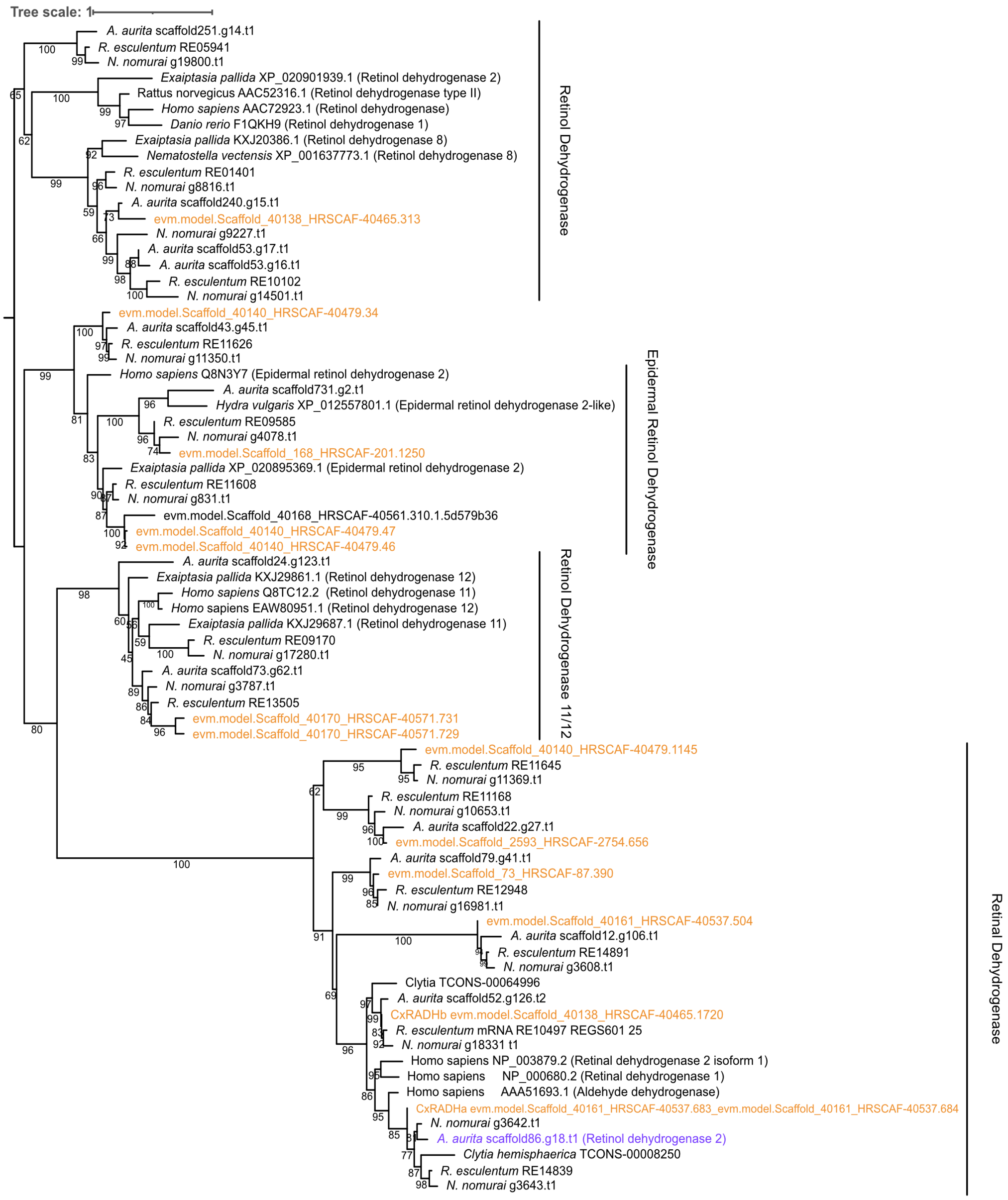


**Figure S2: *C. xamachana* possesses two dehydrogenases closely related to RDH1 and RDH2 of *Aurelia aurita*.**

Phylogenetic reconstruction of cnidarian retinal/retinol dehydrogenase with vertebrate sequences as references. Putative alcohol dehydrogenase genes from four scyphozoan species (*Cassiopea xamachana, Aurelia aurita, Nemopilema nomurai,* and *Rhopilema esculentum*) identified by homology search with BLAST were incorporated into a phylogenetic tree previously published by Poliakov et al. (*31*). *C. xamachana* sequences are highlighted in orange. Short-chain dehydrogenase groups are classified according to the human annotations. *RDH1* and *RDH2* originally described in Fuchs et al. (2014) are indicated in purple. *CxRADHa and CxRADHb* identified in this study are noted with a red asterisk.

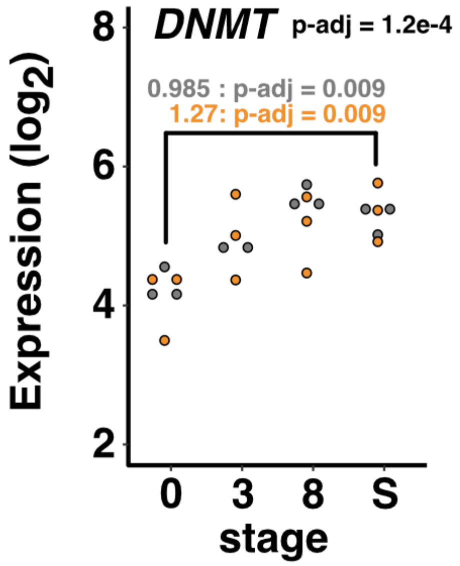


**Fig. S3. Expression of *DNMT* in *C. xamachana* gradually increases as the polyp approaches strobilation.**
*DNMT* gene increases dramatically during strobilation in *Aurelia aurita*. The *de novo methyltransferase* (*DNMT*) in *C. xamachana* gradually increased in expression over the course of two weeks after exposure to symbionts. Read counts were normalized in DESeq2 and log2 transformed. P-adjusted score from the ImpulseDE2 analysis is indicated in black. P-adjusted scores from the two DESeq2 analyses are shown in gray and orange. S= Strobilation.

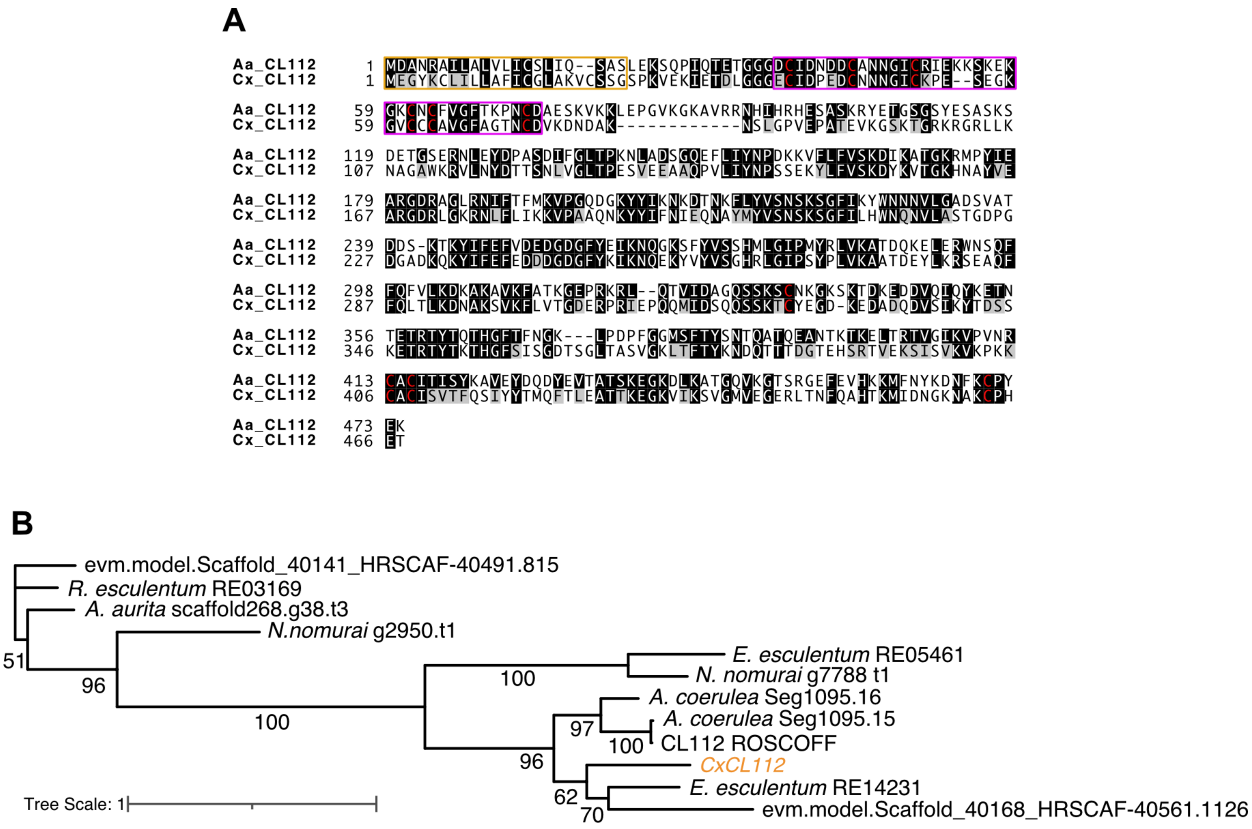


**Figure S4: *Cassiopea xamachana* possesses a sequence with high homology to the CL112 gene from *Aurelia aurita*.**

A) Sequence alignment of *Aurelia aurita* CL112 (AGN03864.1) and *Cassiopea xamachana* CL112 (evm.model.Scaffold_40168_HRSCAF-40561.1132_evm.model.Scaffold_40168_HRSCAF-40561.1133). Alignment was performed using ClustalW and visualized with BoxShade. The two sequences contained 213 identical positions out of 490, and an additional 85 positions with similar chemical properties. The conserved epidermal growth factor-like domain containing six cysteine residues is highlighted with a magenta box. The cysteine residues are indicated in red. The signal peptide sequence is highlighted in an orange box. B) Phylogenetic reconstruction of CL112 within Scyphozoa.

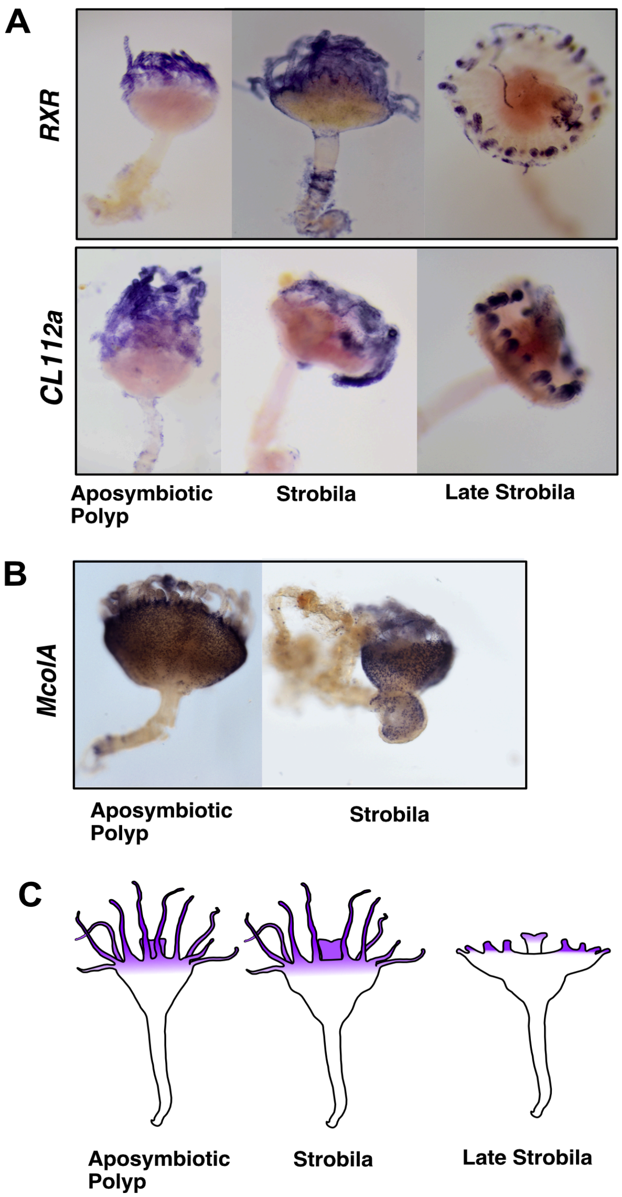


**Figure S5: Whole mount *in situ* hybridization localization of *CxRXR* and *CxCL112* mRNA to the apical region of the polyp.**

Purple staining indicates mRNA localization of *CxRXR* and *CxCL112* (A) in aposymbiotic scyphistomae and strobila. *Minicollagen A* (B) was visualized as a control, which showed the expected "salt-and-pepper" pattern, with staining seen throughout the polyp body and tentacle, but absent from the stalks. Transcripts of both *CxRXR* and *Cx112* were largely localized to the apical portion of aposymbiotic scyphistomae and mid-strobila, but was reduced to the tentacles and hypostome in the late-strobila. (C) Visual summary of mRNA localization of *CxRXR* and *Cx112* during progression of *C. xamachana* strobilation.

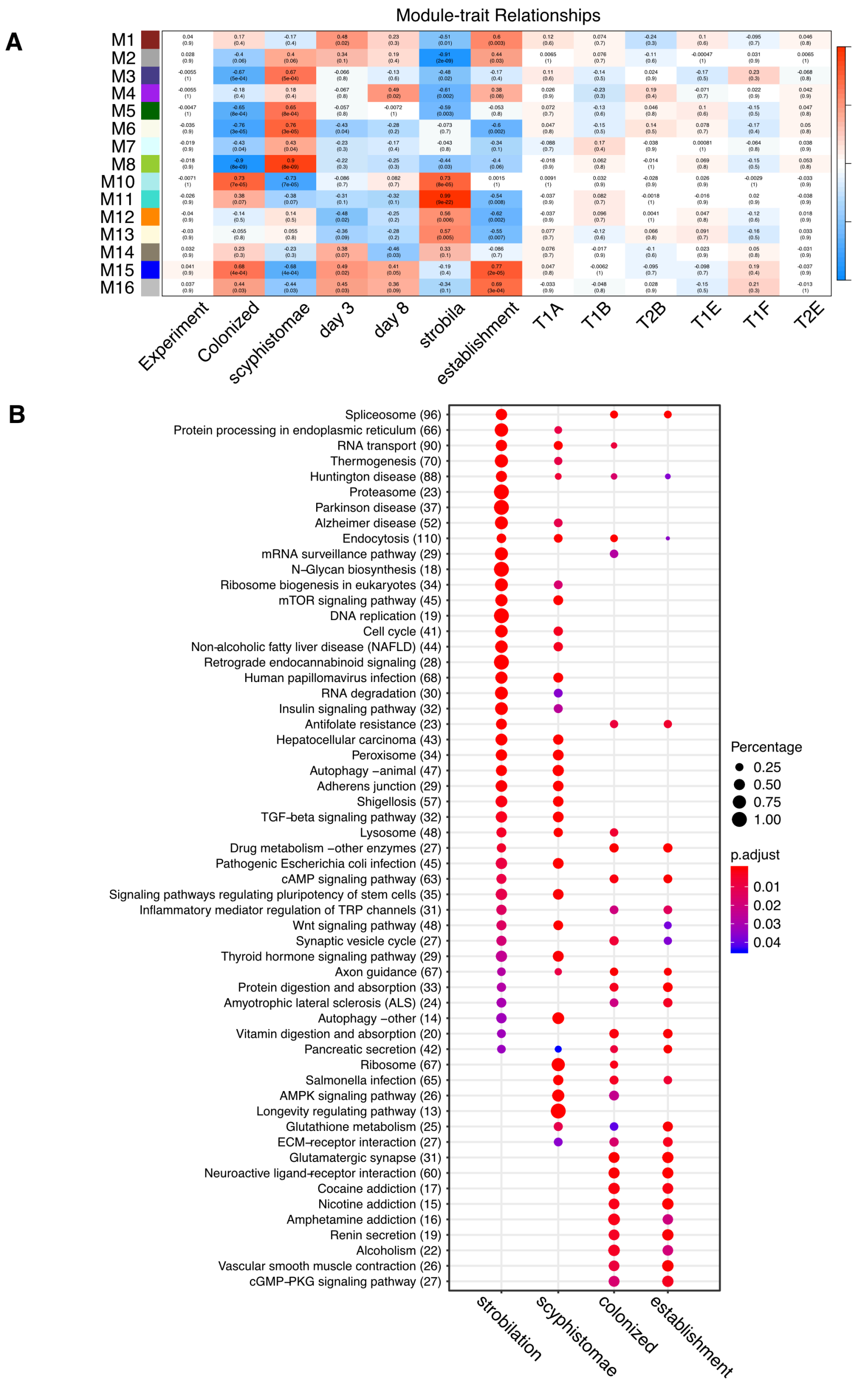


**Figure S6: WGCNA identified gene sets positively correlated with strobilation and onset of symbiosis.**

A. Module eigengene (row) correlation with traits and strains. Each cell contains a correlation value (top) and p-value (bottom). Modules 1, 2, and 14 were significantly positively correlated with establishment of symbiosis (3 and 8 days post-colonization). Modules 9, 10, 11, and 12 were significantly correlated with strobilation (p-value < 0.05). Strength of correlation ranges from -1 to 1, with values above 0 indicating positive correlation. B) Enriched KEGG pathways associated with trait correlated genes.

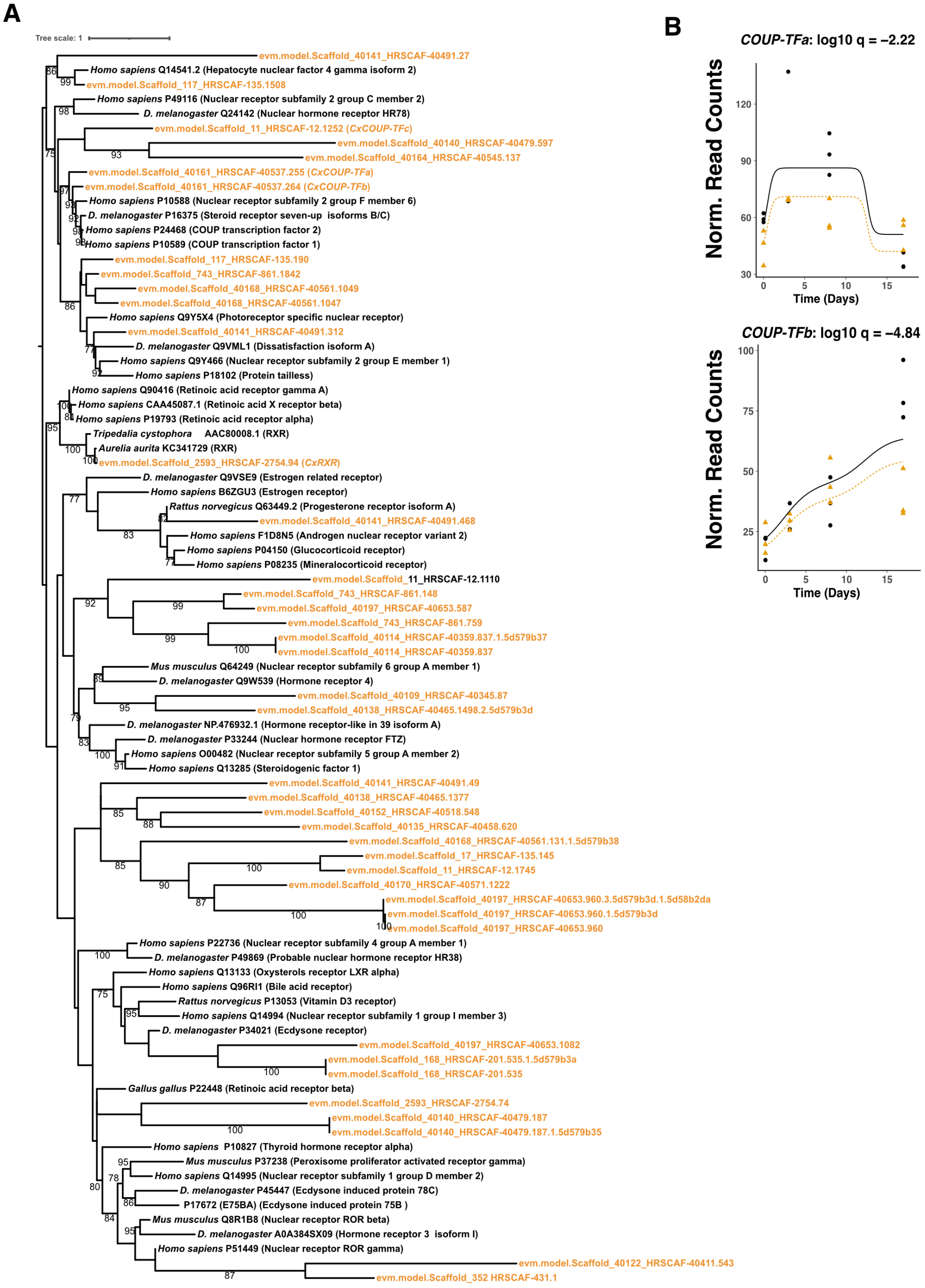


**Figure S7: *C. xamachana* possesses several key nuclear receptors potentially involved in strobilation.**

A. Maximum-likelihood tree of the nuclear receptor (NR) superfamily. *C. xamachana* sequences are highlighted in orange. Representative sequences were retrieved from the NR and UniProt databases. Sequences were aligned with MAFFT and the phylogenetic tree was constructed with iqTree. Bootstrapping = 1000. B) Normalized read counts of *CxCOUP-TFa* and *CxCOUP-TFb*.

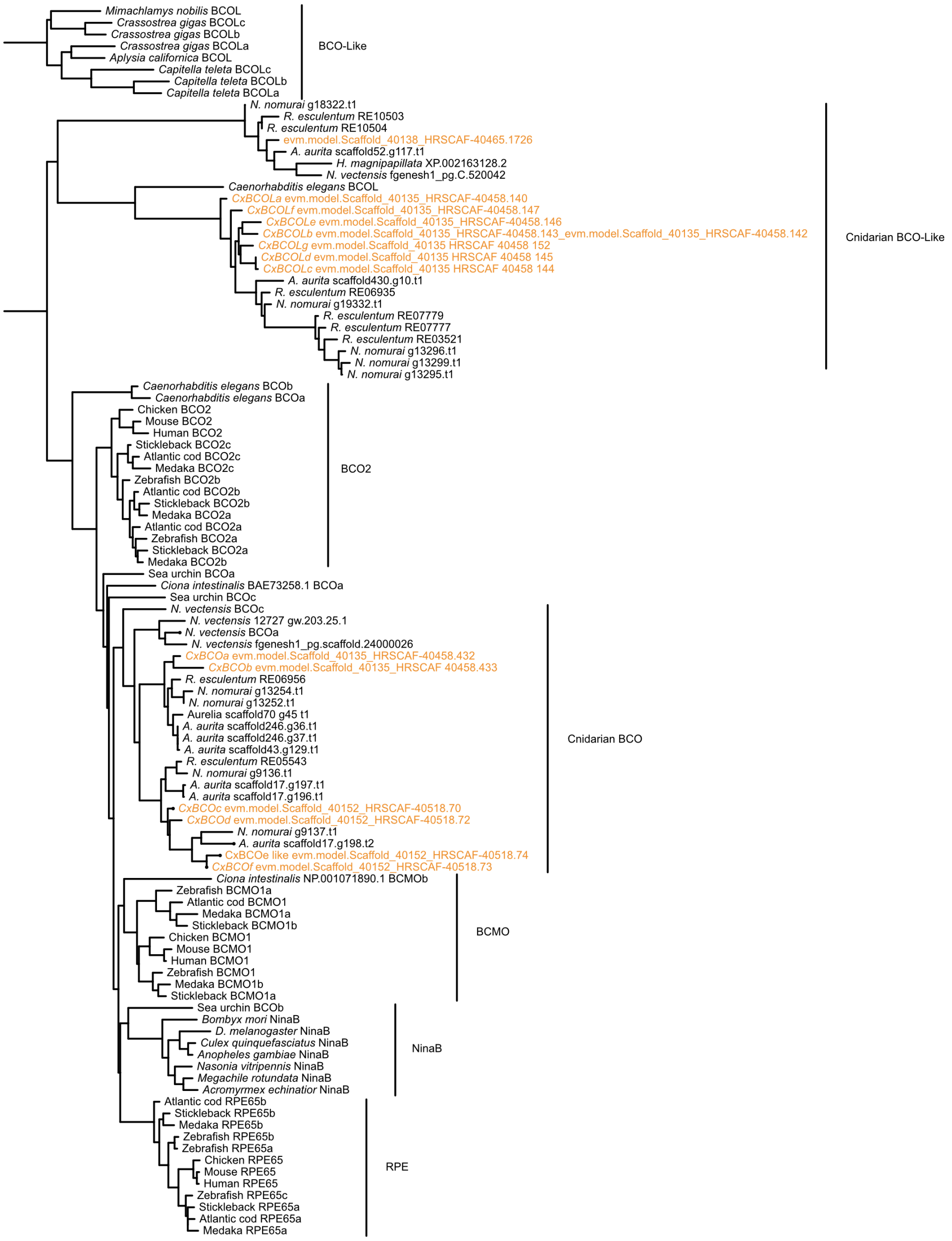


**Figure S8: *C. xamachana* possess multiple orthologs of β-carotene oxygenase and β-carotene oxygenase-like genes.**

*C. xamachana* sequences are highlighted in orange. Representative sequences were retrieved from the nr and UniProt databases based on a previously published tree (*36*). Cnidarian β-carotene oxygenases were largely clustered within one clade, with β-carotene oxygenases-like sequencesclustering separately. Sequences were aligned with MAFFT and the phylogenetic tree was constructed with PhyML. Bootstrapping = 500.

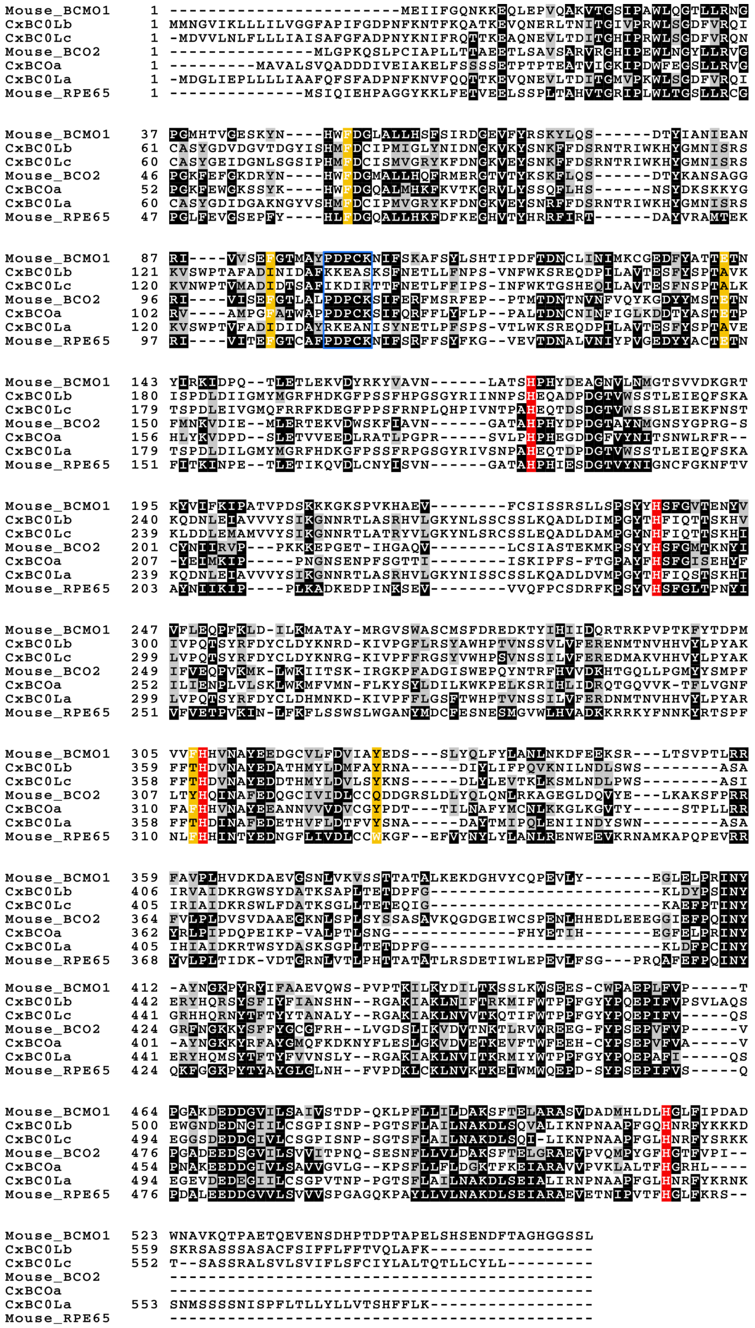

**Figure S9. *C. xamachana* BCO and BCOL genes retain key residues necessary for catalytic activity.** *C. xamachana* BCO and BCOL sequences were compared to representative amino acid sequences from mammalian species. *C. xamachana* sequences were initially identified with BLAST, and confirmed through phylogenetic reconstruction. Amino acid sequences were aligned with ClustalW. Conserved residues important for catalytic activity (red) and substrate binding (yellow) are highlighted. The conserved PDPC(K) motif, necessary for palmitoylation, is conserved in CxBCOa, but lacking in the BCOL genes.

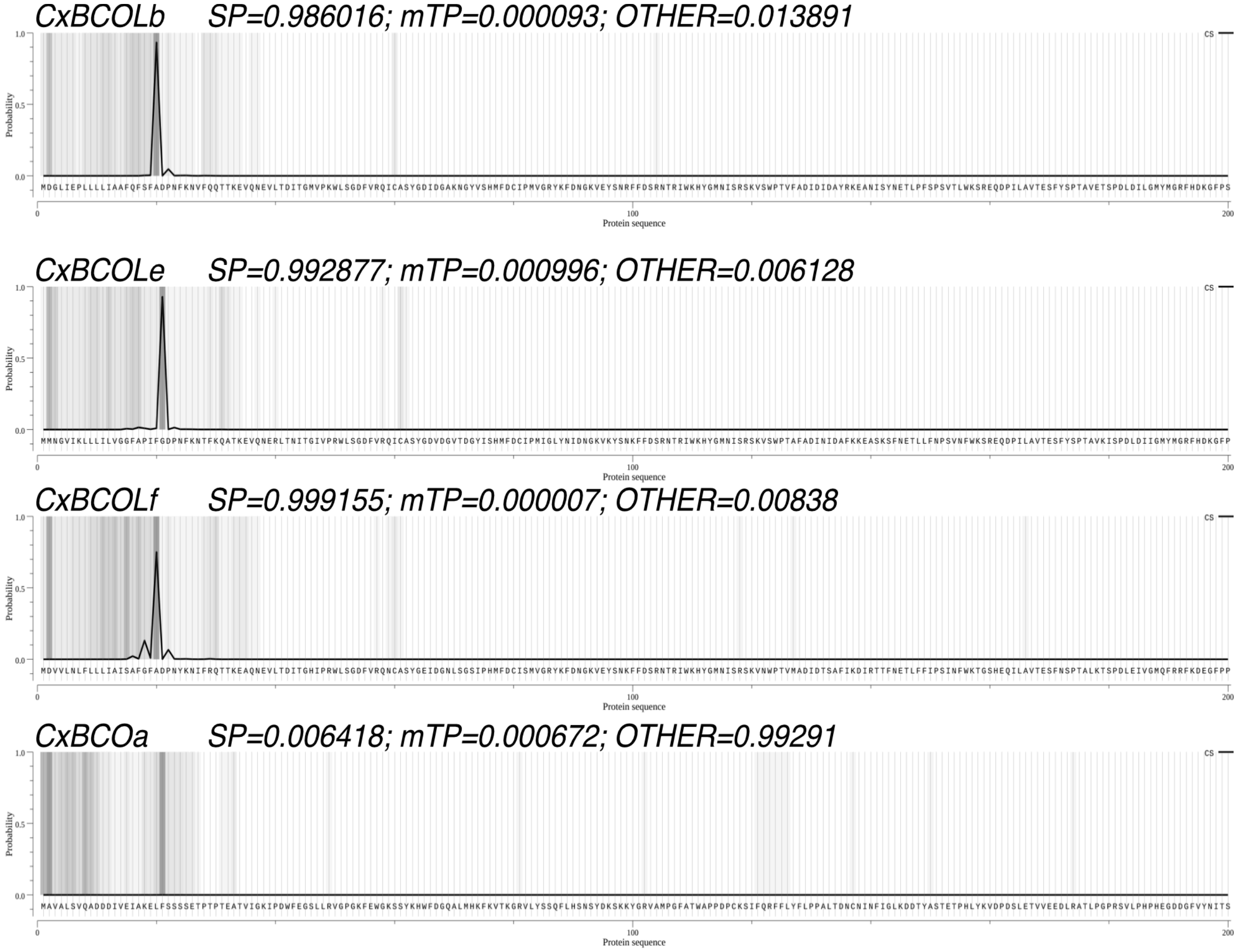


**Fig. S10. SignalP prediction confirms signaling peptides in the N-terminal end of *BCO-like* genes but not BCO.** Presence of signal peptides (SP) and mitochondria transport peptide (mTP) were predicted with TargetP. CxBCOLs were predicted to contain signaling motifs at the N-terminus with high probability, while both SP and mTP were undetected in CxBCOa.

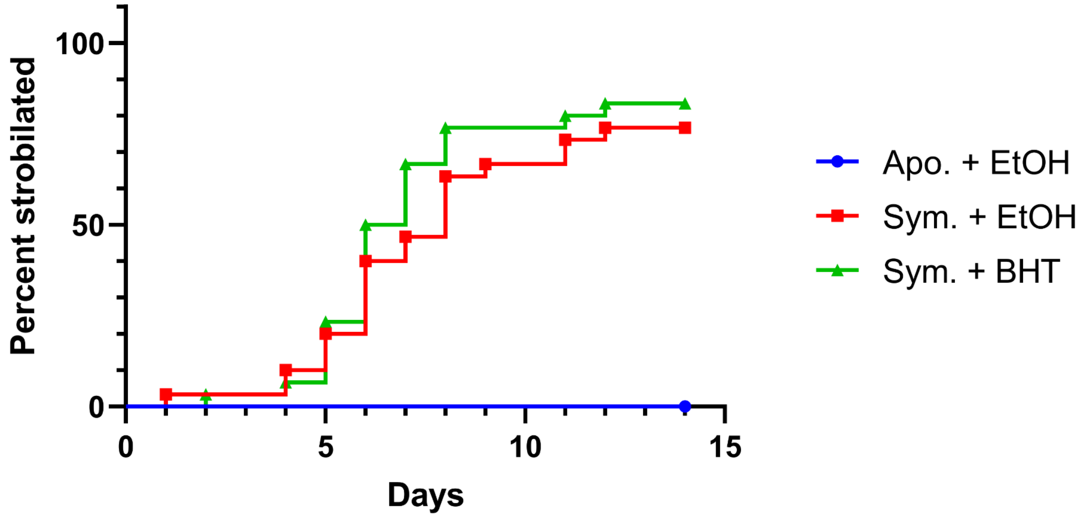


**Fig. S11. Treatment of symbiont polyps with the BCO inhibitor BHT did not delay strobilation.**
Colonized polyps (n=30) were treated with butylated hydroxytoluene (BHT), an inhibitor of the BCO genes. Strobilation rate did not change significantly when animals were treated with BHT.

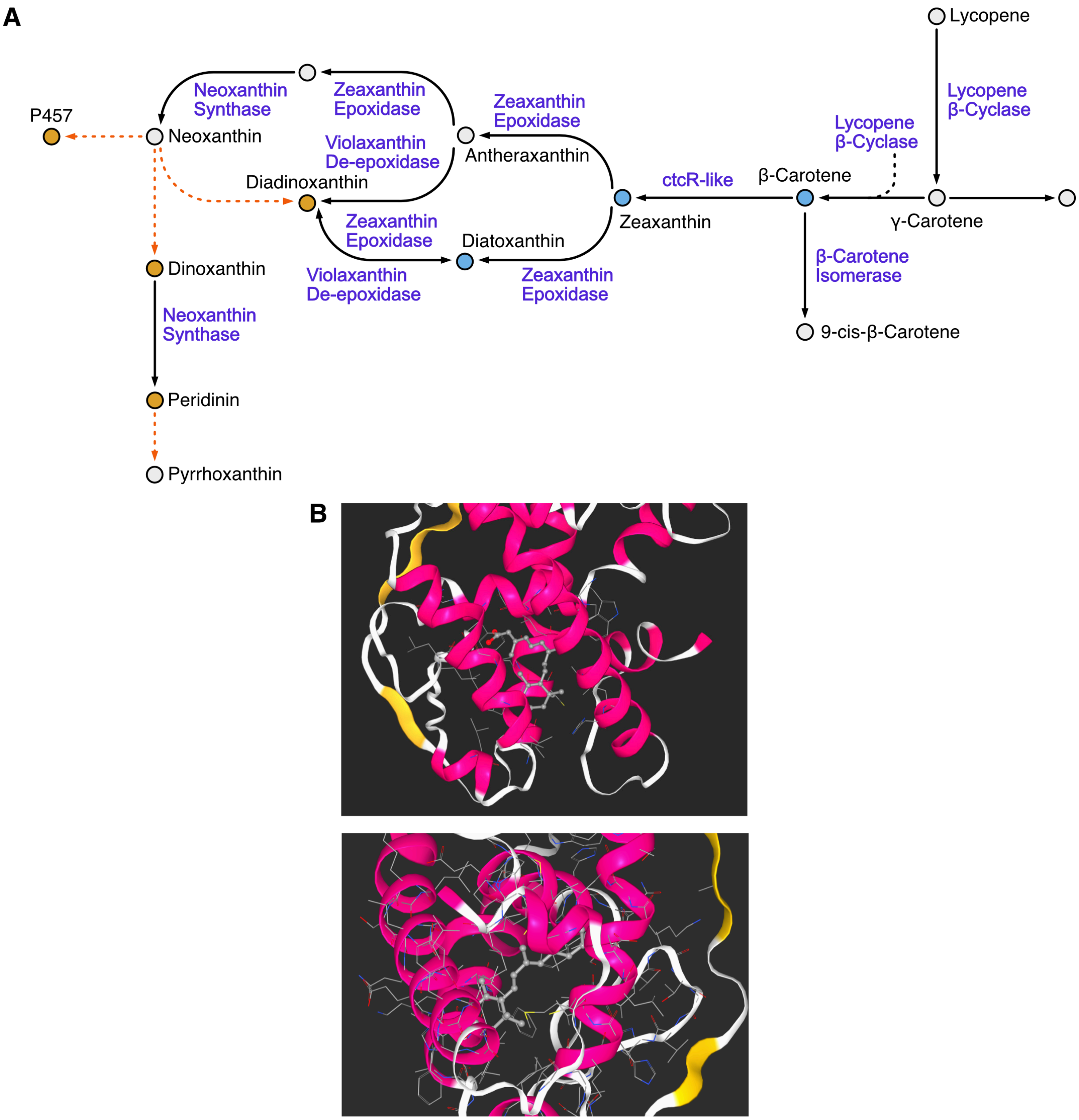


**Fig. S12. The dinoflagellate symbiont is capable of producing precursor carotenoids that can produce potential ligands that can bind CxRXR. A.** Proposed pathway for carotenoid biosynthesis in the dinoflagellate family *Symbiodiniaceae*. All enzymes responsible for catalyzing the reactions (purple) were found to be expressed by *S. microadriaticum* in the RNAseq dataset *(37)*. Carotenoid pigments predominantly found within *Symbiodiniacieae* are shown with orange circles, while low abundance pigments are colored with light-blue (*38, 39*). Grey circles indicate likely intermediates within the biosynthetic pathways. Red dashed lines indicate reactions in which the catalyzing enzyme is yet to be identified. B. 9-cis retinoic acid (A) and zeaxanthin (B) docked within the ligand binding pocket of CxRXR. Ligand binding pocket was predicted with kDeep, and modeling of ligand-protein interaction was determined with bindScope. Both 9-cis retinoic acid and zeaxanthin was calculated to have similar binding affinities to CxRxR (value?).

Table S1. Stats of the *Cassiopea xamachana* genome (version 2.0).

| | |
|---|---|
| Scaffold Number | 735 |
| Total Size (bp) | 366,849,413 |
| N50 (bp) | 17,884,241 |
| L50 | 8 |
| Top 20 Scaffolds (bp) | 363,001,378 |
| # of predicted genes | 29, 645 |
| # of predicted genes w/ isoforms | 32,538 |
| $BUSCO_{COMPLETE}$ | 85.6% (817/954) |
| $BUSCO_{Complete+Fragmented}$ | 93.9% (896/954) |

Table S2. Primer sequences and expected amplicon length for *RXR, CL112,* and *McolA in situ* hybridization RNA probe construction.

| | Forward Primer | Reverse Primer | Length (bp) |
|---|---|---|---|
| RXR | GTGGCACTGTACATTCCTCTGA | AGAGACGAACTGCACTTACCTG | 1057 |
| CL112 | TGCTAGCAATCTCGTCCAGTTT | CAGGTGCATGGAAGAGAGTTCT | 1042 |
| Minicollagen-A | GTGGAATGGGATGTGCTCCTA | GTTGGTGCACATGATGGCATA | 451 |
